# Reduced cerebrovascular reactivity in youth with complex congenital heart defects

**DOI:** 10.1101/2025.11.03.686431

**Authors:** Claudine J. Gauthier, Zacharie Potvin-Jutras, Safa Sanami, Kaitlyn Easson, Guillaume Gilbert, Christine Saint-Martin, Marie Brossard-Racine

## Abstract

**Aims:** Congenital heart defects (CHD) are the most common neonatal malformations. Neonates with complex CHD present with cerebrovascular dysfunction, including deficits in cerebrovascular reactivity (CVR), a measure of vascular reserve. However, it is unknown whether these deficits persist beyond the perioperative period.

**Methods and Results:** Here, we compared CVR between 53 youth with CHD and 54 age-matched controls without CHD. CVR was derived using a novel approach based on resting state blood oxygen-level dependent magnetic resonance imaging. We found that youth with CHD present with relative CVR deficits in the whole gray matter and in the anterior cerebral artery territory when compared to controls. Sex differences were identified in the middle cerebral artery territory, with females having lower relative CVR than males in the CHD group. Greater CVR deficits were observed in individuals with single-ventricle as compared to participants with a two-ventricle physiology. Finally, in the CHD group, a lower CVR was found to be associated with reduced performance on the Metacognitive Abilities Index of the BRIEF-A.

**Conclusion:** These results indicate that cerebrovascular deficits in CHD survivors persistent into young adulthood and that CVR offers great promise as a biomarker of cerebrovascular health which could be targeted in future interventions.

## 1. Introduction

Congenital heart defects (CHD) are the most common neonatal malformations, affecting 1% of births in Canada^1^. To survive, neonates with complex CHD undergo open-heart surgery using cardiopulmonary bypass during the first weeks of life. Improvements in clinical care have greatly increased survival, and more than 90% of neonates with CHD now survive well into adulthood, converting this primarily pediatric condition into an adult chronic condition^2^.

Neonates with CHD present with cerebrovascular dysfunction, as evidenced by reduced cerebral blood flow (CBF) before surgery^3–5^. Previous work has shown that these deficits persist, with female youth with CHD having lower CBF compared to healthy peers^6^. While CBF reflects the amount of blood going to tissue at rest, it is not a measure of vascular reserve, or the ability of blood vessels to meet additional demand. Vascular reserve can be measured as cerebrovascular reactivity (CVR), which reflects vasodilation amplitude in response to CO_2_ inhalation. This is analogous to a cardiac stress test and captures vasodilatory capacity. CVR deficits have been reported in young infants with CHD^7,8^, but no study to date has investigated CVR beyond the first three years of life in this group. This may be partly due to the discomfort and anxiogenic effects of CO_2_ inhalation. However, recent work established that CVR can also be extracted accurately from resting-state functional MRI data^9,10^, making CVR measurements in CHD accessible for the first time.

During childhood, more than 60% of individuals with CHD present with developmental challenges^11–13^ and more than 40% of adolescents and adults with CHD present with cognitive deficits^14–17^. Executive functions (EFs) refer to a group of higher-order cognitive processes such as working memory, cognitive flexibility and inhibition controls. Deficits in EF have been described as the hallmark cognitive challenge experienced by CHD survivors^15,18^. These deficits have been linked to lower academic achievement, quality of life and employment rates in this population^19–21^. While a range of EFs are affected in CHD, metacognitive abilities may be especially vulnerable, especially later in development^21^. Moreover, adults with CHD are at higher risk of cognitive decline as they age^14,22,23^, with a 2.6 increased risk for early dementia^22,23^. Given ongoing work showing that CVR may be one of the earliest detectable biomarkers in the pathophysiological process of dementia^24–27^, especially in populations with vascular risk, CVR may be a valuable biomarker of cognitive decline in CHD survivors.

To address these gaps in our understanding of cerebrovascular health in CHD, this study compared CVR between youth with CHD and healthy peers and assessed the presence of sex-specific effects given our previous findings in this group. We also aimed to explore the functional impact of CVR deficits by evaluating the associations between CVR and EF in individuals with CHD as well as examining their relationships with clinical risk factors.

## 2. Methods

### 2.1 Participants

This cross-sectional study included 53 youth with CHD (25 males & 28 females) and 54 age-matched controls (22 males & 32 females), aged between 16 and 24 years. Individuals with CHD all underwent open-heart surgery using cardiopulmonary bypass before two years of age. Participants were ineligible if they had contraindications for MRI or could not communicate in French or English. Additional exclusion criteria included a history of brain tumors, severe brain anomaly, congenital infections, multisystem dysmorphic conditions, traumatic brain injury, cerebral palsy, or an identified genetic disorder. Written informed consent was obtained from participants aged 18 and older. For those under 18, both the participant and their legal guardians provided written informed consent. This study was approved by the Pediatric Research Ethics Board of the McGill University Health Centre. Individual and clinical information were collected through medical chart review and questionnaires. The family’s socio-economic status was determined using the Hollingshead index^28^.

### 2.2 MRI Acquisition

Whole-brain MRI data were acquired on a 3T Philips Achieva X using a 32-channel head coil. Resting state blood oxygenation level-dependent (BOLD) fMRI data were acquired using a T2*-weighted gradient echo echo planar sequence (repetition time = 2.6 seconds, echo time = 30 ms, resolution = 3 mm isotropic, volumes = 200) for the relative CVR analysis. Additionally, a T1-weighted image was collected using a turbo field echo pulse sequence (repetition time = 8.1 ms, echo time = 2.7 ms, inversion time = 1010 ms, flip angle = 8°, resolution = 1 mm isotropic). The MRIs were reviewed for anomalies by a neuroradiologist, who was blinded to participants’ medical histories.

### 2.3 Image Processing

All images underwent visual quality assessment and only images of sufficient quality underwent pre- and post-processing. The resting-state fMRI BOLD images were preprocessed using Statistical Parametric Mapping (SPM12), FMRIB’s software library (FSL)^29^ and nilearn (*https://nilearn.github.io/stable/index.htm*l). The BOLD images were corrected for motion artifacts using the *Realign* function in SPM12, followed by skull stripping with FSL’s *Brain Extraction Tool*^30^. Slice timing correction was then performed using SPM12’s *Slice Timing* function. Finally, spatial smoothing (FWHM = 6 mm) was performed using *smooth_img* from nilearn. Time domain filtering was omitted from the preprocessing pipeline, as the lower frequency components of the BOLD signal were essential for the relative CVR analysis.

### 2.4 Relative CVR Analysis

Relative CVR maps were measured from preprocessed resting-state BOLD fMRI data^9^. Initially, BOLD images were low-pass filtered (0-0.1164Hz) using *scipy*.*signal*^31^ to extract a proxy for natural cerebral CO_2_ fluctuations^32^. The filtered BOLD signal was then used as a regressor in a general linear model to compute CVR indices, which were subsequently normalized by the whole-brain average CVR. The processing pipeline for relative CVR from preprocessed BOLD data is displayed in ***Figure 1***. Relative CVR maps were registered to MNI-152 space using Advanced Normalization Tools (*https://github.com/ANTsX/ANTs*). A gray matter (GM) mask, segmented from the MNI-152 template using FSL FAST^33^, was applied to the registered relative CVR maps. Because cerebrovascular properties are likely to be determined by vascular territory rather than GM-based anatomical boundaries, a cerebral arterial territory atlas was employed to extract regions of interest (ROIs) in the anterior cerebral artery (ACA), the middle cerebral artery (MCA) and posterior cerebral artery (PCA) territories^34^.

**Figure 1.**
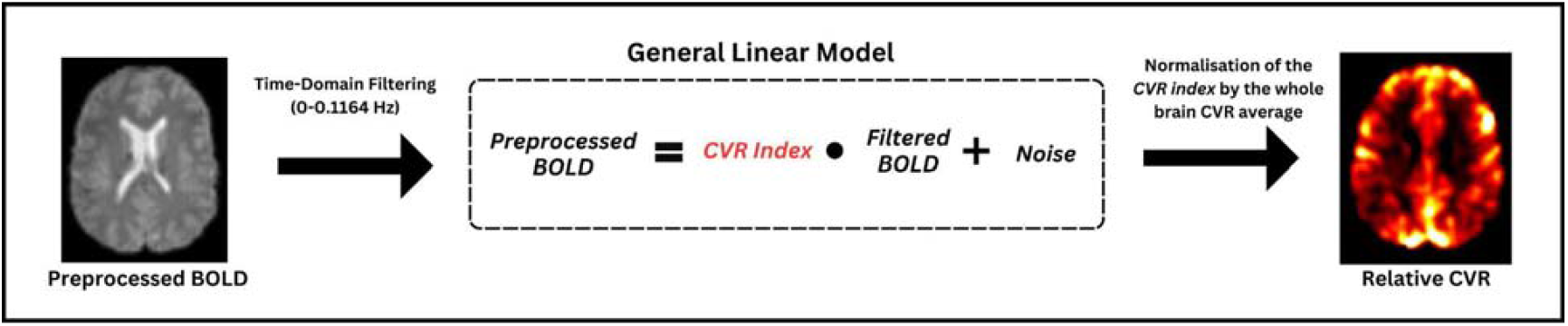
Relative CVR pipeline from preprocessed resting state fMRI BOLD data.

### 2.5 Cognitive Assessment (BRIEF-A)

The Behaviour Rating Inventory of Executive Function - Adult scale (BRIEF-A) is a self-report questionnaire designed to assess EF as manifested in everyday life^35^. The BRIEF-A provides a Global Executive Score, which is the sum of the Behavioural Regulation Index and the Metacognition Index. The specific subscales included in the Behavioural Regulation Index are: inhibition, task shifting, emotional control and self-monitoring. While the Metacognition Index includes initiation, working memory, planning and organization, task monitoring and organization of materials subscales. Scores are reported as T-scores, where higher values reflect poorer executive function^36^.

### 2.6 Statistical Analysis

Normality of the data was assessed using the Shapiro-Wilk test, and relative CVR values were log-transformed to achieve a normal distribution. Group differences in descriptive statistics were evaluated using ANCOVA analyses to compare individuals with CHD and controls, controlling for age and sex for all comparisons except for the age, which was only controlled for sex. These analyses examined differences in education (ordinal), socioeconomic status (SES), and the BRIEF-A summary scales. Sex distribution differences were assessed with a binomial regression, controlling for age. Additionally, group differences in relative CVR were assessed across the whole GM and cerebral arterial territories (ACA, MCA, PCA). Binomial regression was conducted to examine group differences in the sex distribution. To investigate sex differences, ANCOVA analyses were conducted in each group to assess the relationship between sex and relative CVR across the whole GM and ROIs, controlling for age. To assess the effect of severity, we performed ANCOVA analyses to examine the effects of the type of cardiac physiology on relative CVR in individuals with CHD, controlling for age and sex. Additionally, we used multiple linear regression analyses to assess the effects of the number of surgeries on CVR, controlling for age and sex. Lastly, to evaluate the interaction between relative CVR and group on executive function, multiple linear regression analyses were performed controlling for age, sex and SES. Analyses were corrected for multiple comparisons using the false discovery rate (FDR) Benjamini-Hochberg method.

## 3. Results

### 3.1 Participant’s characteristics

Two subjects (one CHD & one control) were excluded due to poor data quality, resulting in a final sample of 52 CHD and 53 controls. The sex specific distribution of cardiac physiology is presented in **Table 1**.

**Table 1.**

| Cardiac physiology characteristics (n=52) |  |  |
| --- | --- | --- |
| Single-Ventricle Physiology (n = 7) |  |  |
|  | Females (n = 4) | Males (n = 3) |
| DILV, n (%) | 1 (14) | 1 (14) |
| Tricuspid Atresia, n (%) | 1 (14) | 1 (14) |
| PA-IVS, n (%) | 1 (14) | 0 (0) |
| HLHS, n (%) | 1 (14) | 1 (14) |
| Double-Ventricle Physiology (n = 45) |  |  |
|  | Females (n = 24) | Males (n = 21) |
| ToF, n (%) | 6 (13) | 7 (16) |
| d-TGA, n (%) | 10 (22) | 8 (18) |
| VSD, n (%) | 2 (4) | 3 (7) |
| Truncus arteriosus type 1, n (%) | 1 (2) | 1 (2) |
| PA-IVS, n (%) | 1 (2) | 1 (2) |
| DORV, n (%) | 2 (4) | 0 (0) |
| TAPVC, n (%) | 2 (4) | 0 (0) |
| COA, n (%) | 0 (0) | 1 (2) |
| Number of Open Heart Surgeries (n = 52) |  |  |
|  | Females (n = 28) | Males (n = 24) |
| 1, n (%) | 17 (33) | 12 (23) |
| 2, n (%) | 6 (12) | 8 (15) |
| 3, n (%) | 4 (8) | 2 (4) |
| 4, n (%) | 1 (2) | 2 (4) |

*CHD = Congenital Heart Defect; ToF = Tetralogy of Fallot; DILV = Double Inlet Left Ventricle; PA-IVS = Pulmonary Atresia with Intact Ventricular Septum; d-TGA = dextro-Transposition of the great arteries; VSD = Ventricular septal defect; DORV = Double outlet right ventricle; TAPVC = Total Anomalous Pulmonary Venous Connection; COA = Coarctation of the Aorta*

**Table 2.**

| Demographic | CHD (n = 52) | Controls (n = 53) | p-values |
| --- | --- | --- | --- |
| Age (years) | 19.96 ± 2.39 | 20.46 ± 2.30 | 0.263 |
| Sex (Females/Males) | 28 / 24 | 32 / 21 | 0.471 |
| Participant's Education (n; 2/3/4/5) | (12/10/22/7) | (2/0/13/38) | < 0.001 |
| Hollingshead SES Index | 40.19 ± 11.37 | 50.19 ± 10.32 | < 0.001 |
| BRIEF-A scores |  |  |  |
| Global Executive (T-score) | 55.47 ± 11.36 | 49.42 ± 8.33 | 0.232 |
| Metacognition Index (T-score) | 55.90 ± 10.64 | 50.25 ± 8.85 | 0.195 |
| Behavioural Regulation Index (T-score) | 54.10 ± 11.98 | 48.73 ± 8.76 | 0.521 |

CHD = Congenital Heart Defect; SES = Socioeconomic Status; Education is coded as 1 = elementary school, 2 = high school, 3 = professional school, 4 = CEGEP/College, 5 = University

Group comparisons of the demographic information and BRIEF-A results are presented in **Table** Three subjects (two participants with CHD and one control) had missing data on the BRIEF-A and were excluded from the analyses. Overall, youth with CHD had significantly lower educational attainment than controls at the time of the study (p < 0.001; n^2^p = 0.363) and lower socioeconomic status (p < 0.001; n^2^p = 0.192), while all other variables, including all BRIEF-A scores, did not differ between groups when taking into account age and sex.

### 3.2 Lower CVR in CHD

Participants with CHD revealed significantly lower relative CVR compared to controls in the whole GM (p = 0.024; n^2^p = 0.050) and the ACA territory (p = 0.030; n^2^p = 0.046) (**Figure 2**). No other significant regional differences were found, and the complete results are presented in **Table S1**.

**Figure 2.**
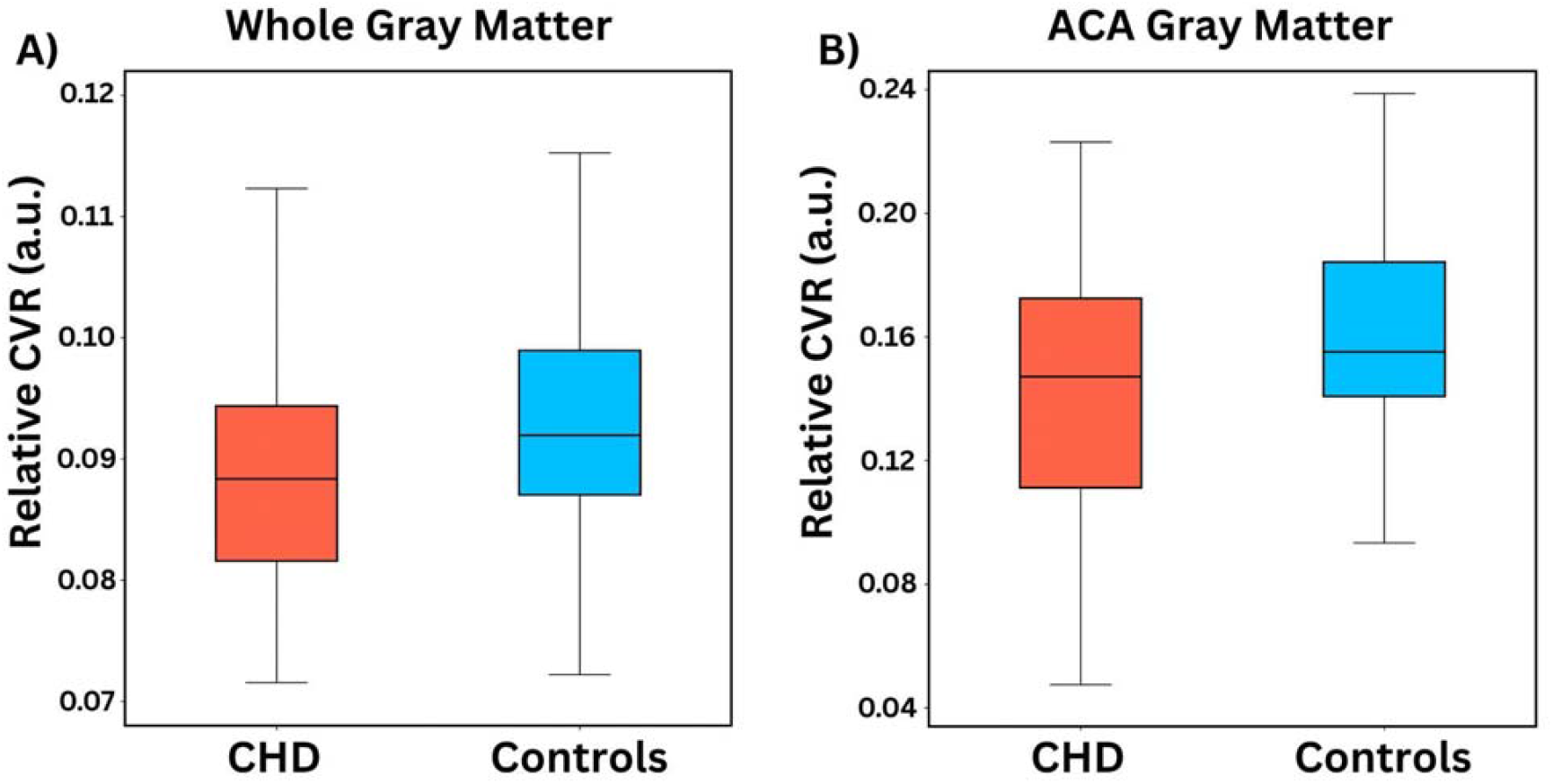
Relative CVR group differences between individuals with CHD and controls. *Box plots from ANCOVA analyses showing the significant relative CVR group differences between individuals with CHD and controls in the A) whole gray matter and B) ACA gray matter territory*.

### 3.3 Lower CVR in females with CHD

In the CHD group, relative CVR was significantly lower in females with CHD compared to males with CHD in the MCA territory (p = 0.005; n^2^p = 0.152) (**Figure 3**). No significant sex differences in relative CVR were found among controls. See **Table S2** for complete results.

**Figure 3.**
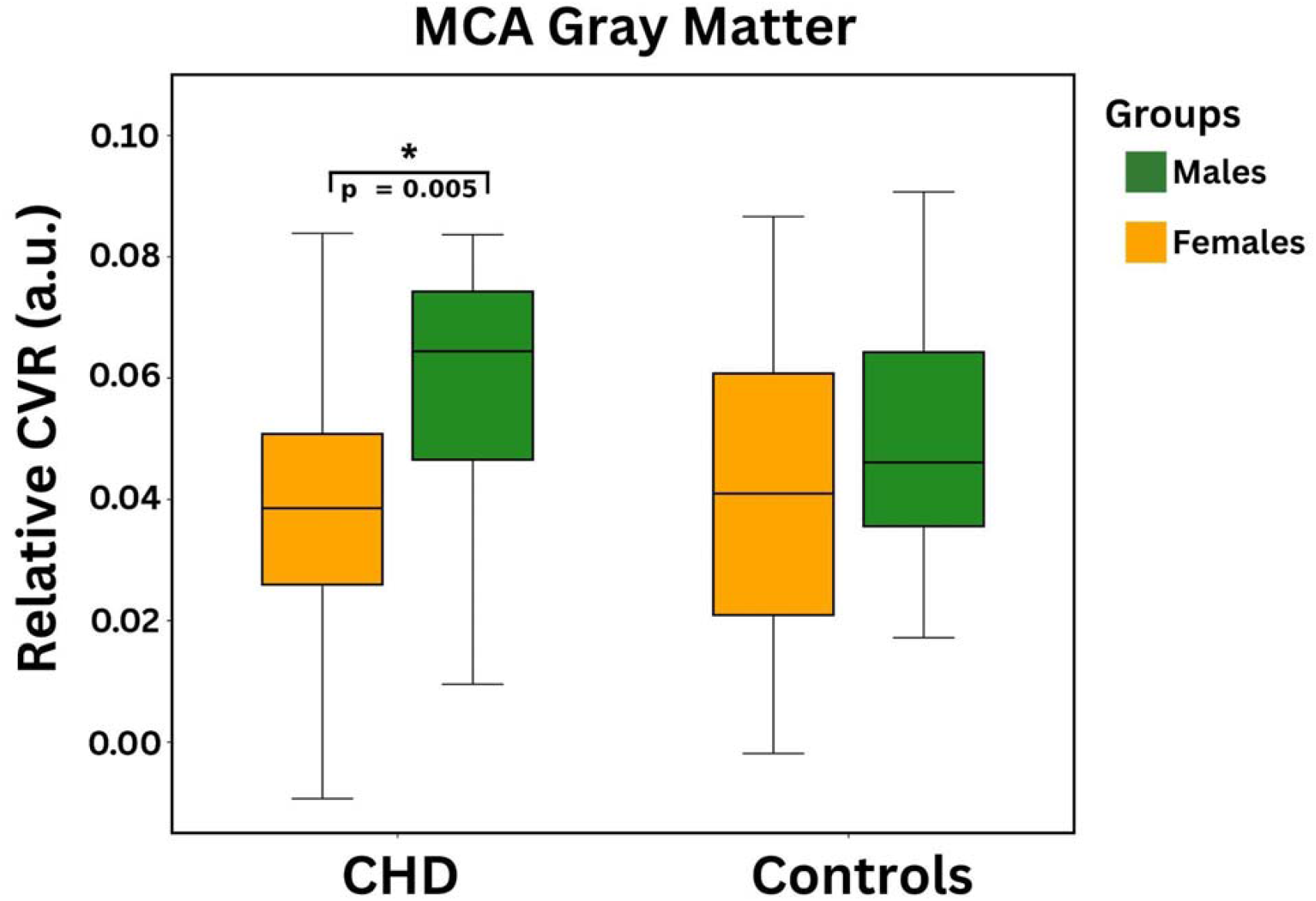
Relative CVR group difference between males and females with CHD. *Box plots from ANCOVA analyses presenting significantly higher relative CVR in males with CHD compared to females with CHD in the MCA GM territory. The p-value shown was FDR corrected*.

### 3.4 Lower CVR in single ventricle physiology

Individuals with a single-ventricle cardiac physiology demonstrated significantly lower relative CVR in the ACA territory (p = 0.006; n^2^p = 0.145) (**Figure 4**) compared to individuals with two-ventricle CHD. No other significant associations were found, and full results are presented in Table S3). Additionally, no significant effects in the relationship between relative CVR and the number of open-heart surgeries were observed (Table S4).

**Figure 4.**
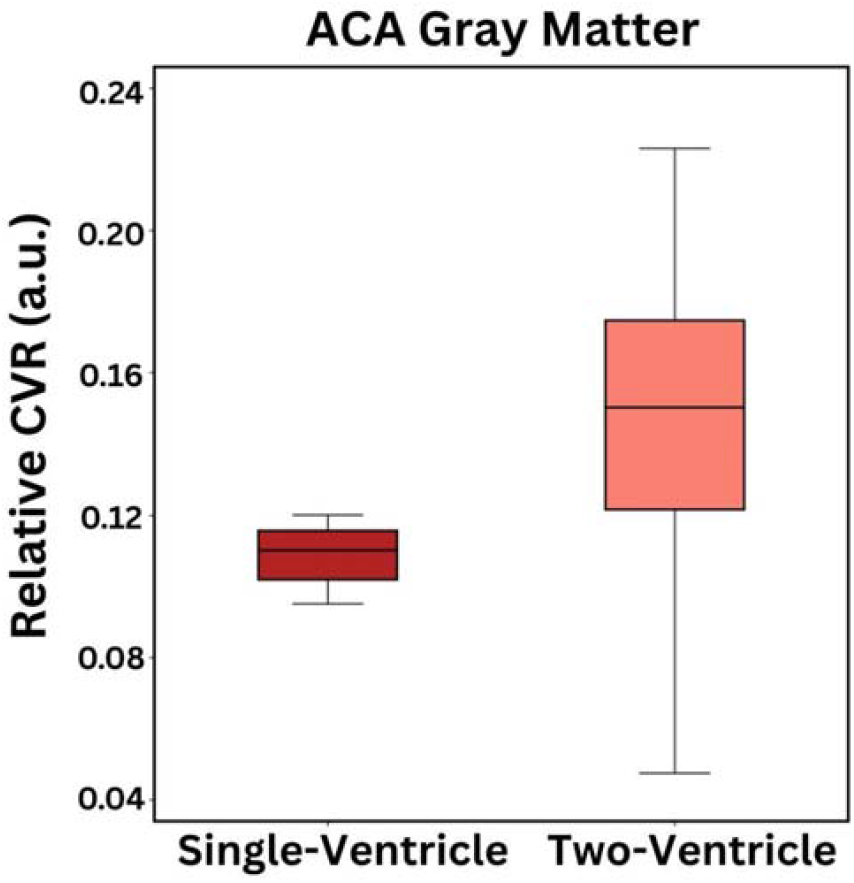
Relative CVR subgroup difference between the type of cardiac physiology. *Box plot from ANCOVA analyses showing the significant relative CVR group difference between individuals with CHD with single ventricle cardiac physiology and with two-ventricle cardiac physiology in the ACA gray matter territory*.

### 3.5 Association between CVR and cognition

The relationship between CVR and cognition was assessed using a linear model including both the overall relationship between CVR and cognitive outcomes, as well as an interaction term to assess slope differences between CHD and control. The overall model was found to be significant for the relationship between whole GM relative CVR and Metacognition Index T-score (R^2^ = 0.076; p = 0.008), as well as a significant group by relative CVR (R^2^ = 0.065; p = 0.007) (**Figure 5**). Furthermore, within this model, we identified a significant group difference in Metacognition Index T-score, showing poorer performance in the CHD group (T-stat = -3.077; p = 0.003). See **Table S5** for the detailed results. In post-hoc disaggregated group analyses, this association between Metacognition Index and whole GM CVR was found to be specific to the CHD group (R^2^ = 0.109; p = 0.025), with no significant relationship in the control group. No significant relationships were found for the Global and Behavioural Regulation indices.

**Figure 5.**
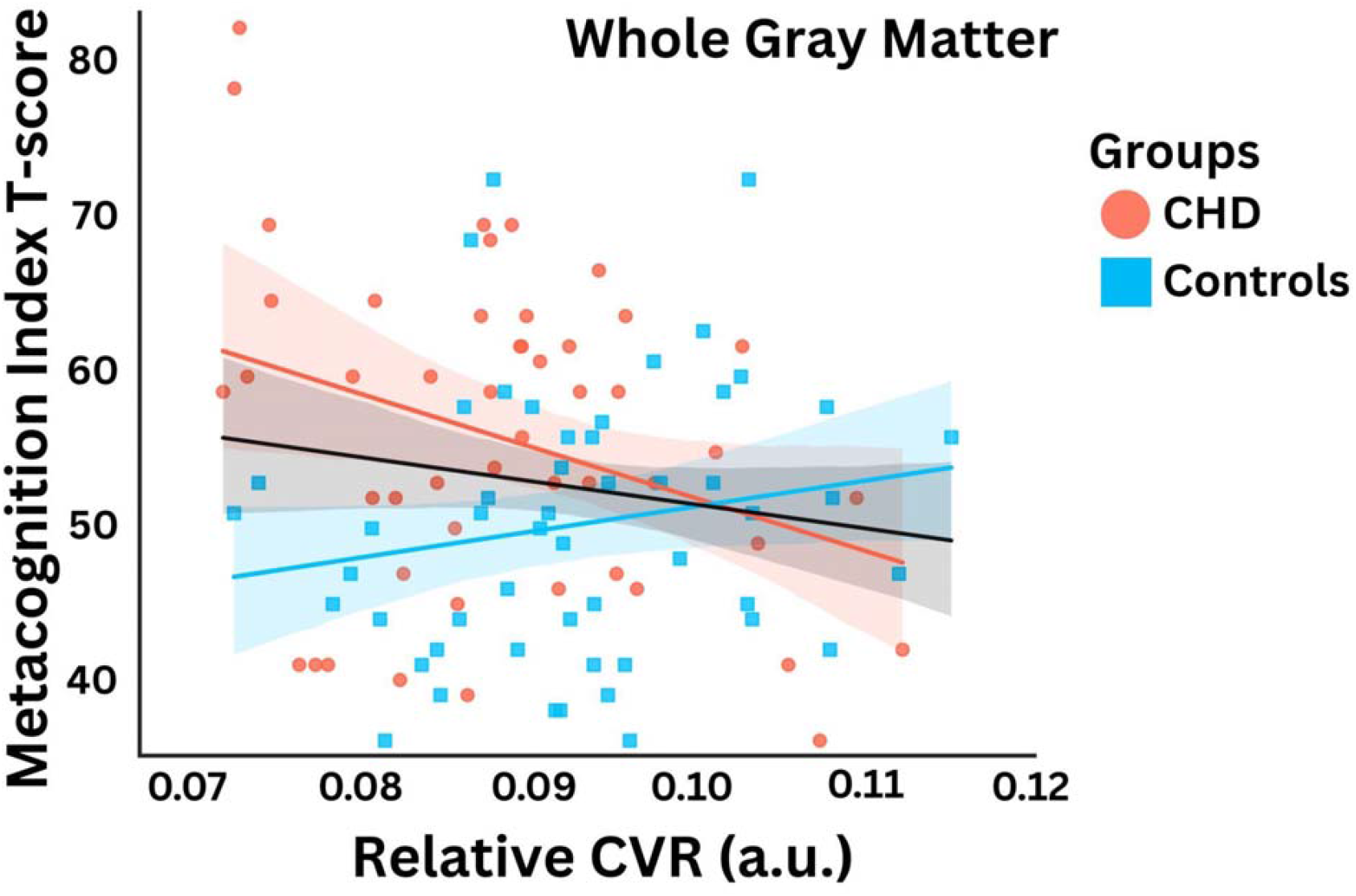
Effects of relative CVR on Metacognition Index. *Multiple linear regression exhibiting a significant group interaction between relative CVR and Metacognition Index T-score in individuals with CHD and controls within the whole gray matter. It is important to note that higher T-score reflects poorer cognitive performance*.

## 4. Discussion

The current study identified CVR deficits in youth with CHD, still present more than a decade after their first open-heart surgery. Importantly, our results demonstrate that while CVR deficits were present in all CHD when compared to controls, females with CHD were found to have lower relative CVR than males in the MCA territory. Furthermore, deficits in the ACA were more pronounced in individuals with a single-ventricle physiology. Finally, lower CVR across the whole GM was found to be associated with poorer executive function in participants with CHD only.

### 4.1 CVR deficits in individuals with CHD

The observed deficit in relative CVR in youth with CHD as compared to controls is well aligned with the previous reports of reduced CVR in neonates and infants with CHD^7^. Although the lack of longitudinal data covering infancy and early adulthood prevents us from concluding on the presence of a direct relationship, our data suggest that some of the cerebrovascular health issues reported in infants persist into adolescence and young adulthood.

Because CVR quantifies vasodilatory capacity, low CVR indicates an impaired ability to increase blood flow to meet additional demand. This increases the likelihood of transient hypoxia during episodes of high metabolic demand. Furthermore, since vasodilation in response to CO_2_ is predominantly caused by endothelial release of nitric oxide^37^, this poor CVR may therefore reflect reduced endothelial health. This is consistent with the extant literature on endothelial function in CHD showing poorer endothelial health across a wide range of different CHD physiologies and types of surgeries^38^.

The relative CVR deficits identified in this study were global, but strongest in the ACA territory. Existing studies in childhood have only documented CVR in MCA^7^, likely because it is more easily accessible than other arteries through the transtemporal acoustic window for ultrasound measurement. Few studies have investigated the ACA^39,40^, and none using a reactivity measurement. However, our observation of differences predominantly in the ACA may be attributable to the ACA’s smaller diameter, as well as the fact that it has more anatomical variations than other major cerebral arteries ^41^. Furthermore, while few studies have investigated the ACA in CHD, one study showed ACA pulsatility to be more related to neurodevelopmental outcomes than MCA pulsatility^39^. In other populations, lower CVR has important functional significance, as exemplified by generalized CVR deficits observed in vascular diseases including coronary artery disease^42^, carotid occlusion^43^ and in both Alzheimer’s disease and mild cognitive impairment^24–26^. Our results demonstrate that adolescents and young adults with CHD show cerebrovascular deficit more commonly observed in diseases of aging and could indicate that CHD may be associated with an accelerated vascular aging phenotype^44,45^. Longitudinal studies are needed to assess whether these CVR deficits are static or increase with age.

### 4.2 Lower CVR in females with CHD

The observed CVR deficits detected in the overall CHD sample were found to be greater in females, who showed lower relative CVR than males with CHD in the MCA territory. The greater CVR deficit in females observed in our CHD cohort mirrors our previous observation of a female-specific CBF deficit in this same cohort, ^6^. In this previous work, we showed that female youth with CHD do not show the higher CBF typically observed in females after puberty as compared to males. Combined with our current observation of low CVR more pronounced in females, these results indicate that female youth with CHD may be at higher risk of transient ischemic episodes. This is in line with data showing that while the overall ischemic stroke incidence in CHD is lower in females than males as in the general population, the incidence rate ratio is higher in females than males with CHD (IRR = 12.12 vs 9.37, respectively)^46^.

The lower CVR in females observed here is especially concerning, given recent work showing that CVR is typically higher in females, at least in older adults^47,48^. However, it is worth noting that there were no significant sex differences in relative CVR in our controls, highlighting the need for more research investigating sex differences in cerebrovascular health. Furthermore, while no study to date has systematically investigated sex differences in cerebral artery morphology in the CHD population, studies in the general population have identified sex differences in MCA hemodynamics, with females typically showing higher velocity and CVR with ultrasound^47^.

### 4.3 Lower CVR in single ventricle physiology

Our results indicate that the CVR deficit observed in CHD is more important in individuals with single-ventricle physiology as compared to individuals with a two-ventricle physiology. While our sample of single-ventricle participants was small, these results are consistent with greater dysfunction in individuals requiring more complex surgeries. We did not observe an effect of the number of surgeries, perhaps due to our limited power at higher numbers of surgeries. Future studies should investigate the effects of surgical and neonatal courses on brain health to better understand the possible relationships with vascular reserve.

### 4.4 Relationship between CVR and cognition

Poorer executive functions are frequently observed in adolescents and young adults with CHD^14– 17^. In the current study, these deficits were apparent as seen by a significantly worse T-score on the Metacognition Index in the CHD group when compared to controls when assessed by multiple linear modelling. Interestingly, the Metacognition Index includes only the BRIEF-A subscales that pertain to the executive functions that are purely related to cognitive processing and excludes the ones dependent on emotional processes^49^. This is in line with the literature in older children with CHD showing greater deficits in metacognitive abilities than behavioural regulation^21^. Furthermore, we observed a negative relationship between relative CVR and the Metacognition Index in the CHD group only. This suggests that in youth with CHD, CVR may be a sensitive biomarker of the brain’s ability to use resources to maintain cognition, though future studies are needed to confirm this relationship using hypercapnic-based CVR and more specific measures of cognitive abilities. This hypothesis is consistent with the accumulating literature showing that higher CVR is associated with better cognitive performance, and especially executive functions, in people with dementia and other heart diseases^26,50,51^. Therefore, our results highlight the promise of CVR as a biomarker of impaired cerebrovascular function in CHD. Future studies should seek to determine whether interventions that improve CVR can also improve EF in youth with CHD.

## 5. Conclusion

This study uncovered that youth with CHD experience persistent cerebrovascular deficits, as shown by lower relative CVR when compared to healthy peers. These deficits are more profound in females and in those with a single-ventricle physiology. Future large-scale longitudinal studies that include prospective data collection of clinical variables are needed to understand the evolution of the heart-brain interconnection across the lifespan. Nevertheless, this study provides foundational evidence that CVR may be a promising biomarker of brain health and function in the youth with CHD warrant of further investigation.

## Data availability

The data that support the findings of this study are available from the corresponding author upon reasonable request and approval from our institutional research ethics board.

## Acknowledgements

The authors thank the participants and their families for their involvement in this study, as well as the research staff and the MRI technologists at the ABCD Research Laboratory and the clinicians at the McGill University Health Centre for their contributions.

