## Supplemental materials for "Reduced cerebrovascular reactivity in youth with complex congenital heart defects"

**Supplementary materials**

**Table S1.** Regional relative CVR group comparisons between participants with CHD and controls.

| **CVR Regions** | **CHD (n = 52)** | **Controls (n = 53)** | **p-values** | **n^2^p** |
| --- | --- | --- | --- | --- |
| **Whole Gray Matter** | 0.089 ± 0.010 | 0.093 ± 0.009 | **0.024*** | 0.050 |
| **ACA territory** | 0.145 ± 0.041 | 0.163 ± 0.039 | **0.030*** | 0.046 |
| MCA territory | 0.049 ± 0.023 | 0.045 ± 0.026 | 0.524 | 0.004 |
| PCA territory | 0.128 ± 0.048 | 0.135 ± 0.044 | 0.535 | 0.004 |

*CVR = cerebrovascular reactivity; CHD = congenital heart disease; ACA = anterior cerebral artery; MCA = middle cerebral artery; PCA = posterior cerebral artery; * = Significant p-value after FDR correction*

**Table S2.** ANCOVA analyses assessing the relationship between sex and relative CVR within groups.

| **Regions** | **CHD Males**  **(n = 24)** | **CHD Females**  **(n = 28)** | **CHD group**  **p-value (n²p)** | **Control Males**  **(n = 21)** | **Control Females**  **(n = 32)** | **Control Group**  **p-value (n²p)** |
| --- | --- | --- | --- | --- | --- | --- |
| Whole GM | 0.090 ± 0.010 | 0.088 ± 0.010 | 0.432 (0.013) | 0.095 ± 0.009 | 0.092 ± 0.010 | 0.277 (0.024) |
| ACA territory | 0.150 ± 0.040 | 0.141 ± 0.043 | 0.479 (0.010) | 0.165 ± 0.038 | 0.162 ± 0.040 | 0.775 (0.002) |
| **MCA territory** | 0.058 ± 0.020 | 0.040 ± 0.023 | **0.005*** (0.152) | 0.049 ± 0.027 | 0.042 ± 0.025 | 0.314 (0.020) |
| PCA territory | 0.117 ± 0.045 | 0.137 ± 0.049 | 0.121 (0.048) | 0.127 ± 0.047 | 0.140 ± 0.042 | 0.300 (0.022) |

*CHD = congenital heart disease; ACA = anterior cerebral artery; MCA = middle cerebral artery; PCA = posterior cerebral artery; CHD group and Control group assesses the effects of sex on relative CVR within subgroup analyses; * = Significant p-value after FDR correction*

**Table S3**. Regional relative CVR group comparisons between participants with single-ventricle cardiac physiology and with two-ventricle cardiac physiology.

| **CVR Regions** | **Single-Ventricle (n = 7)** | **Two-Ventricle (n = 45)** | **p-values** | **n^2^p** |
| --- | --- | --- | --- | --- |
| Whole Gray Matter | 0.082 ± 0.008 | 0.090 ± 0.010 | 0.059 | 0.072 |
| **ACA territory** | 0.107 ± 0.023 | 0.151 ± 0.040 | **0.006*** | 0.145 |
| MCA territory | 0.057 ± 0.021 | 0.047 ± 0.023 | 0.210 | 0.032 |
| PCA territory | 0.117 ± 0.066 | 0.129 ± 0.045 | 0.429 | 0.013 |

*CVR = cerebrovascular reactivity; ACA = anterior cerebral artery; MCA = middle cerebral artery; PCA = posterior cerebral artery; * = Significant p-value after FDR correction*

**Table S4.** Linear regression analyses examining the association between the number of open heart surgeries and relative CVR in participants with CHD.

| **CVR Regions** | **β (Estimate)** | **p-values** | **t-stat** |
| --- | --- | --- | --- |
| Whole Gray Matter | -0.0023 | 0.156 | -1.441 |
| ACA territory | -0.0105 | 0.105 | -1.651 |
| MCA territory | 0.0058 | 0.089 | 1.734 |
| PCA territory | -0.0034 | 0.645 | -0.464 |

*CVR = cerebrovascular reactivity; ACA = anterior cerebral artery; MCA = middle cerebral artery; PCA = posterior cerebral artery;*

**Table S5.** Linear regression analyses evaluating the relationship between group (CHD vs. Controls) x relative CVR interaction and metacognition index T-scores within each CVR region.

| **CVR Region** | **Independent variable** | **β (Estimate)** | **p-values** | **t-stat** |
| --- | --- | --- | --- | --- |
| **Whole Gray Matter** | Group | -55.06 | 0.003 | -3.077 |
|  | Relative CVR | -368.88 | 0.008 | -2.694 |
|  | Group x Relative CVR | 534.40 | **0.007*** | 2.734 |
| ACA territory | Group | -24.60 | 0.002 | -3.168 |
|  | Relative CVR | -69.65 | 0.043 | -2.054 |
|  | Group x Relative CVR | 114.82 | 0.019 | 2.390 |
| MCA territory | Group | -12.28 | 0.007 | -2.744 |
|  | Relative CVR | -116.57 | 0.068 | -1.847 |
|  | Group x Relative CVR | 81.72 | 0.186 | 1.332 |
| PCA territory | Group | -13.52 | 0.029 | -2.217 |
|  | Relative CVR | -18.55 | 0.541 | -0.613 |
|  | Group x Relative CVR | 49.69 | 0.254 | 1.148 |

*CVR = cerebrovascular reactivity; ACA = anterior cerebral artery; MCA = middle cerebral artery; PCA = posterior cerebral artery; * = Significant p-value after FDR correction*
